# Assessing motile cilia coverage and beat frequency in mammalian in-vitro cell culture tissues

**DOI:** 10.1101/2023.03.22.533861

**Authors:** Ricardo Fradique, Erika Causa, Clara Delahousse, Jurij Kotar, Laetitia Pinte, Ludovic Vallier, Marta Vila-Gonzalez, Pietro Cicuta

## Abstract

Cilia density, distribution and beating frequency are important parameters lung tissues, for example in diagnostics of Primary Ciliary Dyskinesia, and in the study of *in vitro* models, e.g. derived from induced Pluripotent Stem Cells. Video microscopy can be used to characterise these parameters, but most tools available at the moment are limited in the type of information they can provide, usually only describing the ciliary beat frequency of very small areas, while requiring human intervention and training for their use. We propose a novel and open source method to fully characterise cilia beating frequency and motile cilia coverage in an automated fashion without user intervention. We demonstrate the ability to differentiate between different coverage densities, identifying even small patches of cilia in a larger field of view, and to fully characterise the cilia beating frequency of all moving areas. We also show that the method can be used to combine multiple fields of view to better describe a sample without relying on small pre-selected regions of interest. This is released with a simple graphical user interface for file handling, enabling a full analysis of individual fields of view in a few minutes on a typical personal computer.

## 1. Introduction

The pseudostratified airway epithelium observed *in vivo* is often recapitulated *in vitro* by culturing human airway epithelial cells (hAECs) at the air-liquid interface (ALI). This configuration closely resembles *in vivo* conditions and drives differentiation towards a mucociliary phenotype. The tissue can be started from primary cells or cell lines. Moreover, in the last decade multiple protocols have demonstrated the ability to derive hAECs from induced Pluripotent Stem Cells (iPSCs) [1,2].

*In vitro* models of human airways epithelium are an effective tool to study mucociliary clearance (MCC) and disease pathogenesis in the respiratory tract. However, depending on the cell origin, culture medium and growth factors employed, these cultured tissues may differ for the amount and type of cells (i.e. ciliated, basal, secretory cells). Therefore, number, distribution and ciliary length, and physical properties of the airway surface liquid (ASL) are often highly diverse among the samples, leading to a variability in cilia beating frequency (CBF), coordination and alignment. Moreover, the ability of motile cilia to generate an effective MCC can be altered in the presence of genetic diseases such as cystic Fibrosis (CF), where the ASL hydration level decreases drastically, or primary ciliary dyskinesia (PCD), where cilia have defective function [3]. Hence, the need to measure coverage in motile cilia and CBF in *in vitro* mucociliated tissues in order to quantify physical differences between different cells and different culture protocols and be able to correlate them to the MCC efficiency.

While improvements have been made in recent years, there are still a limited set of tools for the analysis of microscopy videos of ciliated cells, with the standard being the commercial software Sisson-Ammons Video Analysis (SAVA; Ammons Engineering, http://www.ammonsengineering.com/SAVA/sava.html) [4]. Even though new alternatives have appeared in recent times [5], this method typically only measures the CBF and mostly in user selected areas. This usually requires human intervention in either the areas selection or the frequency range filtering, a potential source of bias in the analysis. In addition, and unlike the method presented in this manuscript, SAVA software does not evaluate how much of the visualised area is covered with beating cilia, and thus only allows for CBF comparison of small pre-selected areas. Alternative techniques have been developed over the years, spanning from methods that make use of cheaper cell phone cameras for image acquisition [6], to others attempting to eliminate or automate the Region of Interest (ROI) selection [7], or trying to determine spatial distribution of CBF [8]. However, these techniques are still limited to CBF measurements only, and mostly limited to small areas of the epithelium or only able to run on video of cilia imaged from the side. A different method must thus be developed to better quantify the differences in beating cilia coverage and frequency, particularly in heterogeneous samples that can require the acquisition of a high number of field of view (FOV) to obtain a representative characterization, and for samples that can present the full range from no visible moving cilia to full coverage in motile cilia. Identifying regions with cilia beating is also a pre-requisite for other analysis, for example multiDDM [9] aiming to measure the spatial coherence of ciliary dynamics.

The approach presented here can, with no input from the user, accurately detect areas containing beating cilia, calculate the CBF distribution across the entire FOV and, given a sufficient number of FOV for a particular sample, provide a description of the average percentage of tissue covered in motile cilia, and aggregate to a CBF distribution over an entire sample of FOV from matching conditions, as illustrated in Figure 1.

**Figure 1.**
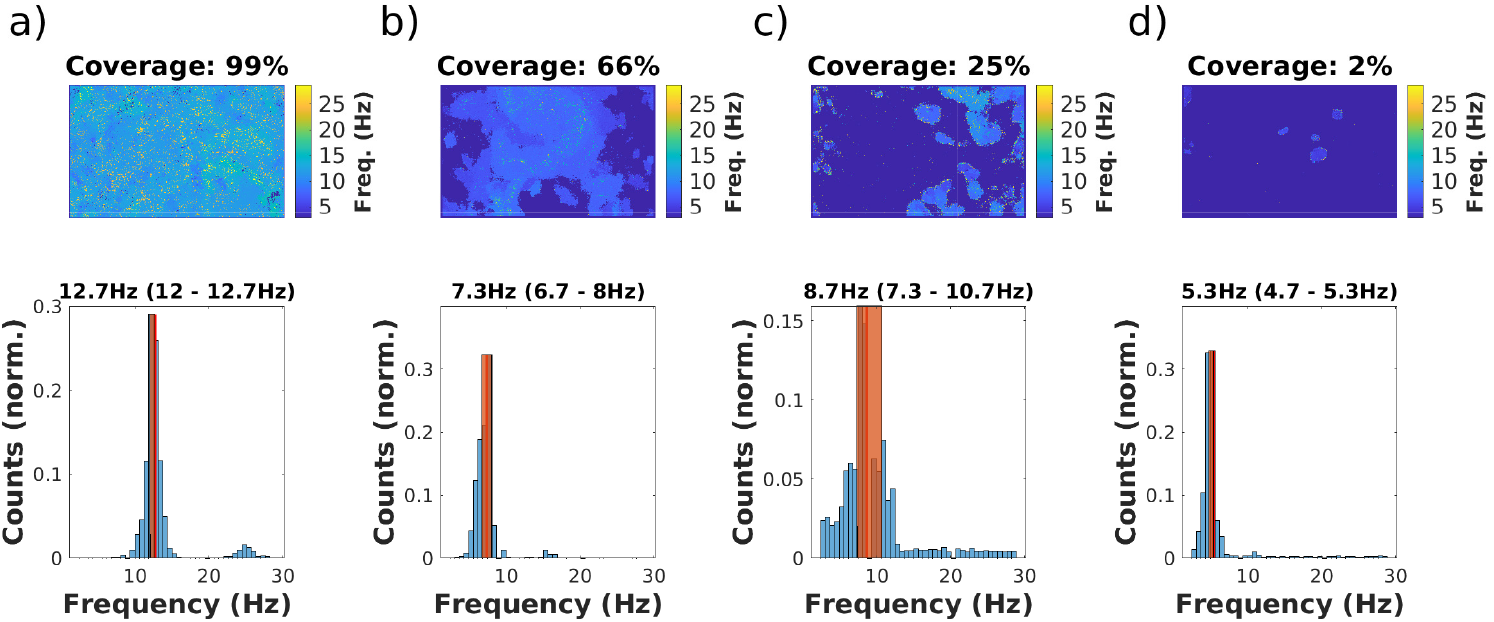
The method presented in this paper allows to determine the areas of an ALI culture where cilia are beating, in conditions of high (a), medium (b) and low (c) coverages, and even a single cilia patch (d). Then distributions of the CBF can be obtained. Each FOV is *≈* 187 µm by 117 µm).

## 2. Ciliated epithelia

The algorithm developed here has been tested on videos collected from *in vitro* airway epithelium derived from both human primary cells and iPSC. Commercially available human Bronchial Epithelia reconstituted *in vitro*(MucilAir™) were purchased from Epithelix Sàrl and maintained following the protocol provided by the company. Other samples have been derived from human iPSC by Laetitia Pinte and Marta Vila Gonzalez at the Wellcome-MRC Cambridge Stem Cell Institute. In both cases, epithelial tissues are reconstituted on a microporous filter in an plastic insert of 6.5 mm diameter and imaged to assess cilia properties at the ALI stage.

## 3. Microscopy setup

For these measurements, digital high-speed videos are typically recorded at least at ∼ 80 frames per second (fps). Our setup consists of a Nikon Ti-E inverted microscope equipped with a 60 × water immersion Nikon objective (NA 1.2), using a CMOS camera (model No. GS3-U3-23S6M-C; Point Grey Research/FLIR Integrated Imaging Solutions, resulting in 1 pixel = 0.098 µm. Each FOV *≈* 187 µm by 117 µm). Samples of ciliated cells are imaged in a custom-made chamber which regulates temperature, CO_2_ and humidity continuously, maintaining values of 37 °C, 5%, and 100%, respectively. The permeable membrane of the insert is kept on a thin layer of culture medium during the experiments.

## 4. Software Development

One of the basic parameters used to evaluate ciliary function and MCC efficiency in airway epithelial tissues is the CBF. The majority of algorithms employed for CBF measurement are based on fast Fourier transform (FFT). Briefly, CBF can be extracted from the frequency of light intensity fluctuation by Fourier transforming the pixel intensity over time, therefore measuring the ciliary cycles. A common limitation of these methods is that to determine the relevant frequency of the pixels across the FOV, it is required to distinguish between areas where cilia are beating and areas without cilia movement. Only the first group should contribute to the final CBF measurement. This might seem like a simple task, but it is in fact hard to develop a robust and automated method working reliably in the general range of possible conditions (very low to high coverage). It is therefore common for these techniques to require human intervention in the selection of the areas to analyse, creating a potential source of bias and limiting the number of FOV that can be analysed in a given time.

For this purpose, an algorithm was developed relying on the differences in frequency and intensity in the Power Spectrum Density (PSD) of areas with and without beating cilia, over different length scales, to first determine the areas where movement is present, followed by a Fourier power spectrum analysis to determine the CBF of the ciliated areas. The first approach identifies areas where beating cilia are likely to be located by observing how the power spectrum changes between regions with and without movement. Areas with no movement are typically characterized by a Fourier Power Spectrum with higher amplitudes in the lowest frequencies, mostly representing the background noise inherent to the camera and acquisition settings. On the other hand, areas with high cilia coverage will present higher and sharper peaks for relatively bigger frequencies.

The development and video analysis was performed using MATLAB R2022b (MathWorks Inc, Natick, Massachusetts), running on a Linux workstation (AMD Ryzen 3900X, 32 Gb RAM, Nvidia RTX 3090 24 Gb).

### (a) Initial image normalization

As the illumination quality may vary across the different videos analysed, and in order to achieve consistency in the gray-scale range among them, each video is first normalized. The normalized intensity at height *i*, width *j* and time *t*(*I*_*n*_(*i, j, t*)), is obtained by subtracting the minimum value measured in the entire video (min(*I*)) to the original intensity signal *I*(*i, j, t*), and dividing the result by the maximum intensity value measured max(*I*):

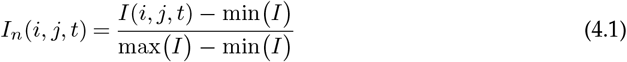

This operation sets the value of all pixels in the video in a range between 0 and 1. Background intensity is then decreased by calculating the average intensity across the frame-stack, and subtracting it from each pixel obtaining 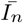.

### (b) Pixel Fourier transform analysis

The frame-stack auto correlation is then calculated, and this processed frame-stack is subsequently used in the FFT analysis. Pixel PSD *P*(*i, j, f*) is estimated for each position (*i, j*) through FFT over time, applying a Hann window to the signal 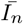. This windowing function touches zero at both ends eliminating all discontinuity at the boundaries of the frequency vector. The resulting spectrum is filtered through a band pass filter (2 Hz to 30 Hz).

### (c) Box average function

Areas with beating cilia within the FOV are determined by observing how each pixel power spectrum changes within square ‘boxes’ of increasing size, centered in each pixel. 15 different box sizes between 5 and 33 pixel are first determined:

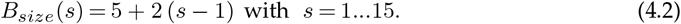

Then, a ‘Box Average Function’ is applied to the original spectrum for each box size. That is, the spectrum for a given pixel is calculated as the average of the spectra of the *B*_*size*_ surrounding pixels, for the different box sizes. A visualization of the key steps in the process is described in Figure 2.

**Figure 2.**
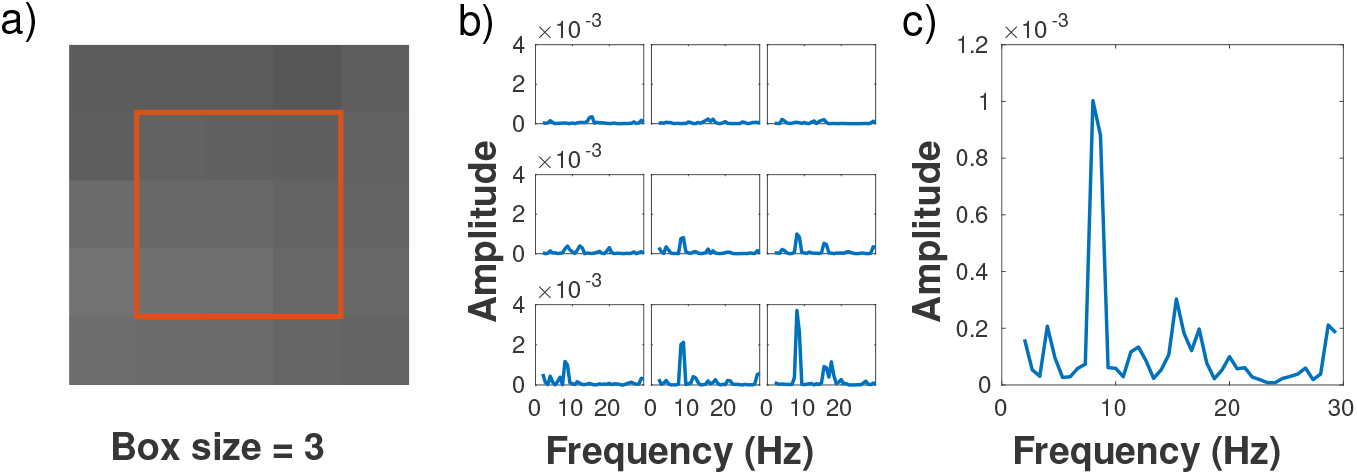
Visual representation of the Box Average function. a) Area selection for a *B*_*size*_ = 3. b) Spectra (of the autocorrelation function) for each pixel in the selected area. c) Final Box Averaged spectrum.

Formally, let the input be a 3D array, *P*(*i, j, f*), with dimensions *H* × *W* × *F*, where each layer in *f* corresponds to the amplitude of the PSD for all pixels in a FOV with *i* height and *j* width for a given frequency *f*. Let the output be a smoothed version of this stack, *A*_*s*_. The box average function is applied by dividing each slice in the stack into overlapping blocks, or “boxes” of size *B*_*size*_ × *B*_*size*_. The output for each pixel *A*_*s*_(*i, j, f*) is then calculated by taking the average amplitude of all the pixels within the box centered in (*i, j*), for each layer *f* in the stack.

This can be expressed mathematically as follows:

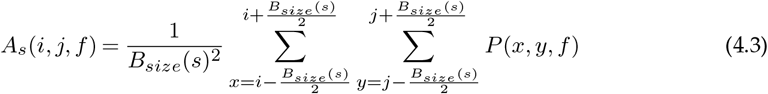

Where *i* = 1….*H, j* = 1…*W, s* = 1…15, *f ∈* [2, 30] Hz.

### (d) Movement map determination

The resulting 3D array is then simplified by fitting each spectrum with a linear least square fit and storing the slope of the fit (see Figure 3 for a visual reference of the overall sequence of steps). Let the input be the previously computed 3D matrix, *A*_*s*_, and let the output be a matrix of the slopes of the best linear fit for the amplitudes along the *f* axis of *A*_*s*_, indicated as the matrix *a*_*s*_(*i, j*). Let *F* be the length of *f*. In this case, for each *s* we apply a linear least square fit to each element of the matrix *A*_*s*_ along its 3rd dimension, using *f* as the independent variable.

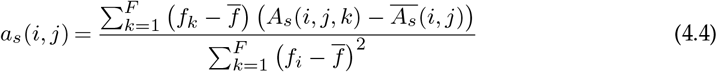

**Figure 3.**
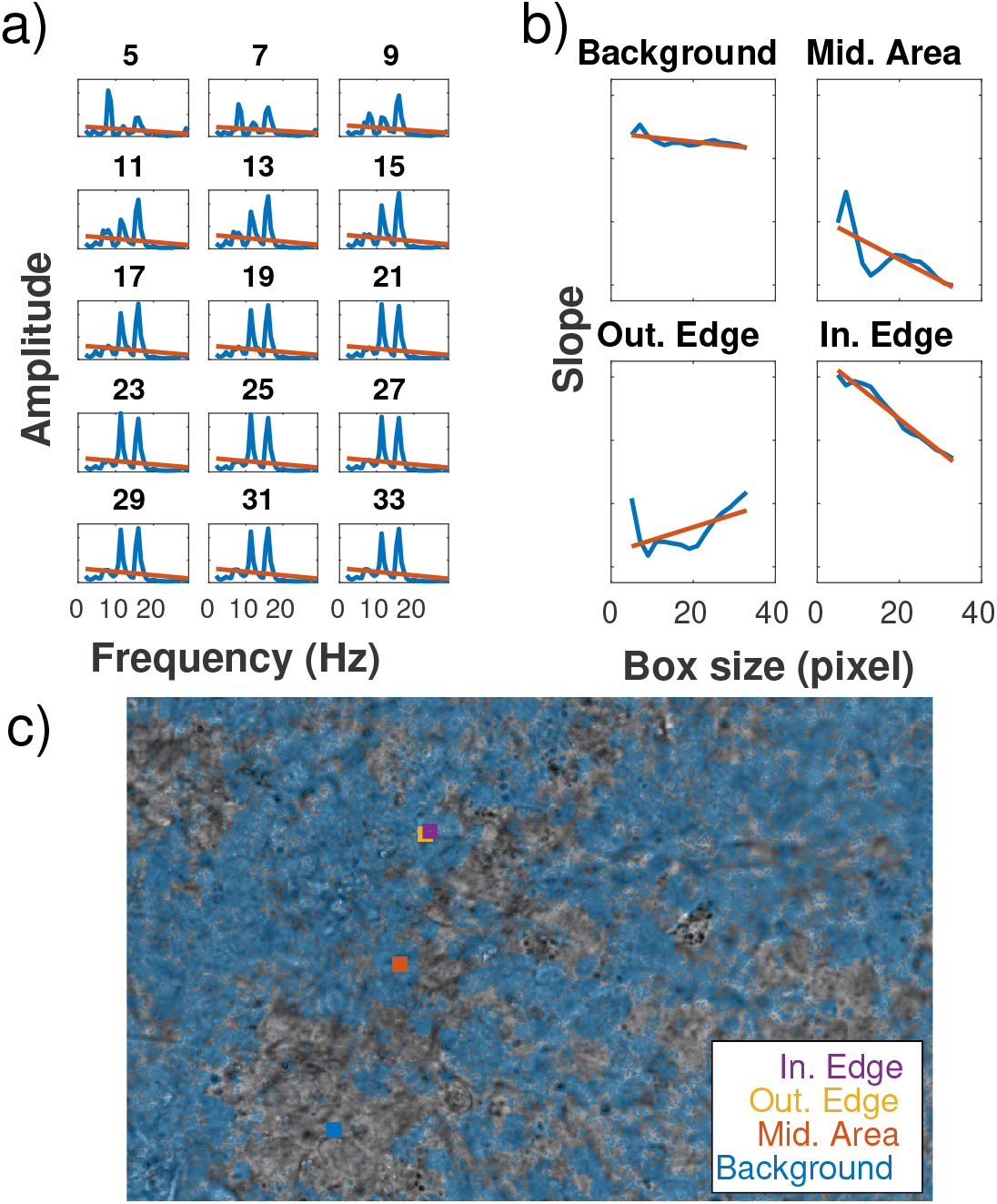
Analysis of frequency spectra, over progressively larger ‘boxes’, leads to identifying four classes of pixels: Those deep inside or outside a ciliated area, and these just outside or inside an edge between ciliated and non-ciliated. Figure shows the processing stages to determine the Movement Map, i.e. the pixels in ciliated regions. a) Example of the slope map *M*_*s*_ obtained for each box size of an individual pixel. b) Estimated variation of *M*_*s*_ over each box size, for the selected areas from 5 by 5 to 33 by 33 pixels, as marked. c) Binarised Movement Map overlaid on top of the FOV, blue shaded areas are marked as presenting movement. Video available in Supplementary Materials and in data repository. Overlaid are selected areas for visualization in the FOV, corresponding to an area just outside of an area containing moving cilia (Out. Edge), just inside of a moving region (In. Edge), in the middle of larger area of movement (Mid. Area), and on an area with no noticeable movement in its surrounding (Background). FOV is *≈* 187 µm by 117 µm).

Where:

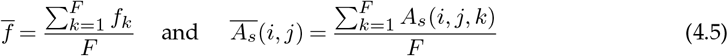

Note that with this linear fit (Figure 3(a)) we are not trying to achieve a good interpolation of the data; the purpose of the fit is to identify in a numerically rapid and robust fashion if there is an outstanding low frequency peak that rises above background or noise. This information is obtained in a particular pixel (*i, j*) for each box sizes, resulting in 15 different 2D matrices *a*_*s*_, *s* = 1 15. The matrices are stacked together is a 3D matrix *M*(*i, j, s*) of size (H×W×15), i.e. an array of “box slopes” (see Figure 3.a for a visual representation of this step, for the case of a single pixel).

A second linear least squares fit is then applied over the box size axis *s*, again storing the value of the calculated slopes, resulting in a final matrix describing the profile changes over the different box sizes for each pixel spectrum in the entire FOV, *m*(*i, j*). Pixels in a background region, that is, away from any movement area, should have relatively stable spectra profiles over the different box sizes, thus low *m* values. Pixels within a high movement area display slightly higher variations with different box sizes due to small local variations in cilia beating frequency. On the other hand, pixels on the edge between a high movement area and a background area should present a decreasing or increasing slope profile, depending on whether they are within the movement area or just outside of it (Figure 3.b and c). The resulting matrix *m* is again box-averaged with a 5 pixel box size to remove noise, and a binary threshold is applied, so that any absolute value under 0.6 × 10^*−*7^ is considered background. This value was determined experimentally as the most flexible regarding FOV with high, medium and low coverage, as well as unfocused FOV, in order to present the least amount of artifacts (Figure 3.d). The obtained movement map is later used to calculate the cilia coverage as the percentage of area containing movement within the FOV.

### (e) Cilia Beating Frequency

To determine the distribution of frequencies within the FOV, the original pixel PSD map *P* is firstly box averaged with a 3 pixel box size to remove noise. The characteristic frequency for each pixel is then determined by calculating the local maxima of the PSD for each box, and selecting the one presenting the highest amplitude (Figure 4.a), generating a frequency map of the FOV (Figure 4.b). The resulting matrix is again filtered to remove any artifact resulting from the box average operations before, by removing an edge of pixels on the outside of the FOV correspondent to the size of the largest box used, and is then combined with the previously determined movement map creating a frequency map for the areas containing movement (Figure 4.c). The characteristic frequency for the FOV is calculated as the median frequency measured, and the corresponding 25th and 75th percentiles are also evaluated to quantify the scatter of the frequencies in the FOV.(Figure 4.d).

**Figure 4.**
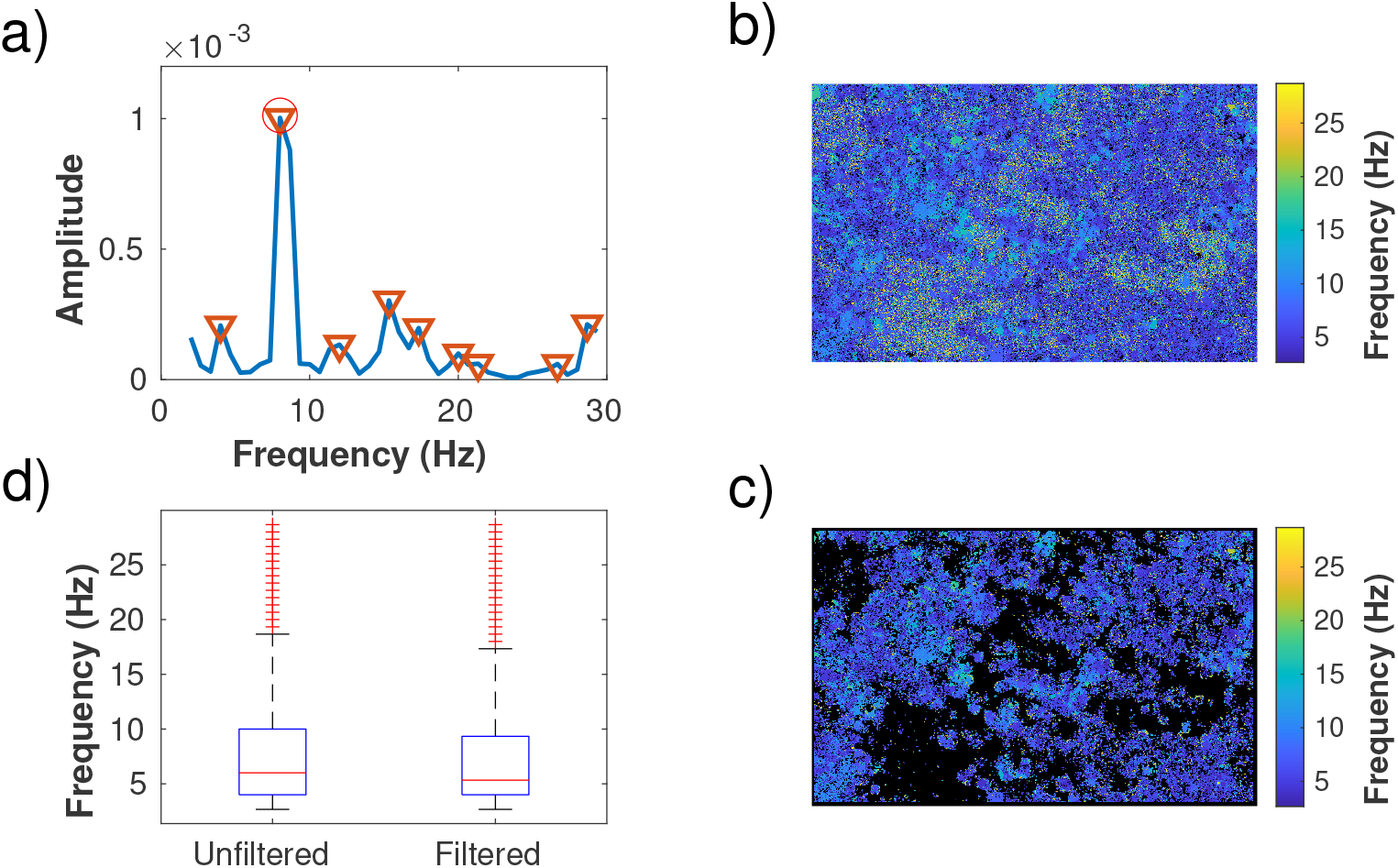
The process of identifying the Movement Map allows a rigorous determination of the CBF distribution in each FOV. a) Example of the characterization of power spectrum from a single pixel, average of the box sizes. Local maxima are marked by (▽); (°) marks the selected frequency. b) Pre-filtering frequency map for the entire FOV. c) Final frequency map, after movement map thresholding; the black areas represent removed pixels. d) Frequency distributions for the FOV of the panels (b) and (c), showing the tighter distribution of CBF following the Movement Map filtering. Making this distribution more robust is one of the outcomes of this analyis; the other is to reliably identify fractions of tissue covered by motile cilia. The FOV is *≈* 187 µm by 117 µm).

In this way it is possible to determine the beating cilia coverage for FOV with different coverage, from low to high, with minimal error, even when only small patches of cilia are detected.

### (f) Detecting focusing issues and debris

As already discussed, *in vitro* cultured hAEC tissues can significantly differ from each other for coverage, CBF, and ASL properties depending on the cell type and the culture technique. Moreover, even following a well-defined imaging protocol, the collected videos may vary for the focusing choices or the presence of blurred elements and debris. For this reason we implemented the algorithm with a debris detection and a bad focus warning. Both these sections are based on each pixel intensity power spectrum and standard deviation data.

#### (i) Bad focus detection

The cells lining the tract of the human airways going from the trachea to respiratory bronchioles form a pseudostrafied epithelium. Here, even though all cells make contact with the basal membrane creating a single cell layer, the nuclei are disposed on different levels, causing the illusion of cellular stratification. In some samples this effect is more pronounced, making it challenging to have all the ciliated cells entirely in focus in the same focal plane. Moreover, the presence of mucus may hinder the correct focusing on cilia when this layer is extremely viscous, leading to a globally blurred image (Figure 5). It is possible to attempt to identify a blurred image by its STD distribution. For each video the STD is computed as:

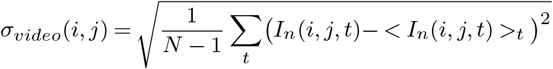

obtaining a *H* × *W* matrix in which each element assumes the value of the pixel standard deviation over time. The computed STD values are normalised to a range between 0 and 1, and the matrix is box averaged with a box size of 10 pixel. The STD distribution in the FOV is then fitted with a log-logistic regression, and the resulting mean (*µ*) and spread (*σ*) are calculated. Being the STD histogram quite sharp when the video is blurred, it is possible to implement a bad focus detection section based on the spread of the distribution. If *σ <* 0.05, a warning message is triggered, and a flag is added into the output file, so that the output provided can be further verified for accuracy.

**Figure 5.**
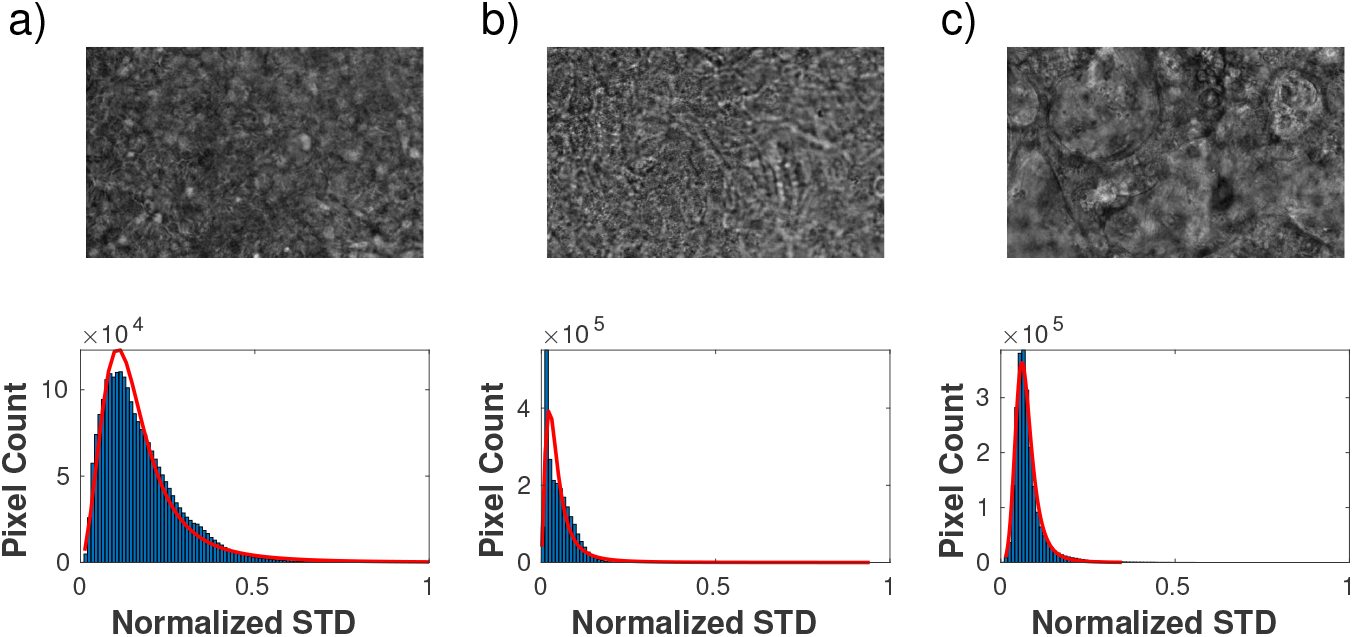
STD histogram distributions for focused (a) and unfocused (b and c) FOV. The difference in profile allows for a rough identification of blurred areas within the FOV. Each FOV is *≈* 187 µm by 117 µm).

#### (ii) Debris detection

It is not uncommon for videos of biological samples to present unwanted debris crossing the FOV. These debris can be detached cells, mucus vesicles or dust particles. While the analysis method described here appears to be robust enough that the presence of small debris should not significantly affect the final result, it still takes note of cases where debris are detectable. The PSD of the pixels crossed by debris when transported across the FOV can present more noise, resulting in less sharp peaks. This can be quantified by measuring the prominence of the highest peak in the PSD (i.e. how much the peak stands out from the surrounding baseline of the signal due to its height and its position compared to other peaks). If a local maxima is part of a range of peaks (i.e. noisy PSD representing background), it is more likely to have a smaller prominence compared to an isolated, sharp one, which commonly describes cilia beating. These information can be collected in a ratio map *R*(*i, j*), where every entry represents the ratio between the prominence and the amplitude of the highest local maxima (both a direct output from the CBF calculation in the previous section). Once computed, the matrix *R* is box averaged with a 10 pixel box size. The entries in *R* which value is close to 1 are more likely to represent areas with beating cilia, due to the presence of a sharp peak in the PSD. On the other hand, if the value of the ratio is small, the pixel presents a noisy spectrum and it is likely to represent an area with debris moving or background. In addition, the presence of a debris can result in an increase in the STD for the pixels in that specific region, due to the variation in the intensity signal compared to the average. The STD map *σ*_*video*_(*i, j*) is calculated and box averaged as described above.

Since the pixels potentially affected by the debris movement are characterised by a high STD signal (moving high contrast feature) and a small ratio value (noisy PSD), a debris mask can be finally calculated as:

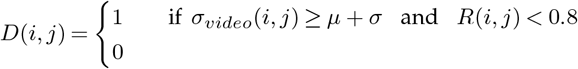

Entries true in the mask would represent pixels possibly crossed by a debris while the video has been captured. If the total number of pixels tagged in the debris mask is higher than 2% of the total pixel count, the FOV is tagged as potentially containing debris (Figure 6).

**Figure 6.**
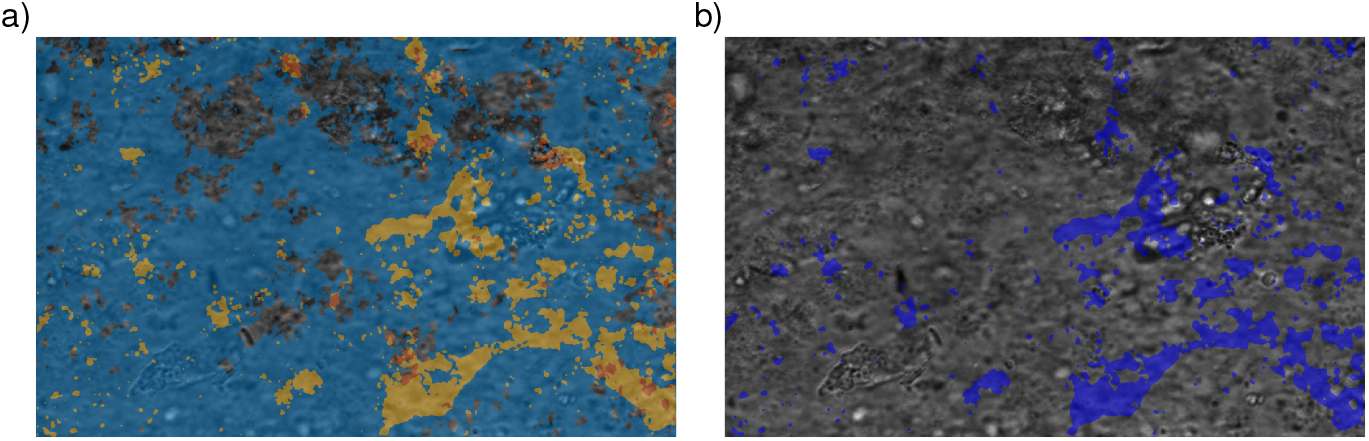
Example FOV containing debris. a) FOV overlay with the areas with value intensity above the threshold for STD in blue, and areas with high ratio values in orange. b) Resulting debris map overlaid on the FOV. Each FOV is *≈* 187 µm by 117 µm).

#### (g) Sample data aggregation

To fully characterise a sample while avoiding any potential bias, it is important to collect several FOV across the entire tissue and aggregate their information. This algorithm includes an aggregation function that can calculate the frequency distribution and average CBF for the entire sample. This is done by pooling together the pixel information relative to each video as one larger FOV describing the whole sample.

#### (h) Graphical User Interface

To make it easier for the user to run the analysis on multiple samples and FOV, it is useful to work through a graphical user interface. This allows the user to input the folder where the data is stored, the output folder where the results should be written to, as well as filename based sample identification and renaming (Figure 7). This input data is saved as a configuration file in the output folder so that in the case of any unexpected shutdown the analysis can continue without the need to re-input all the information.

**Figure 7.**
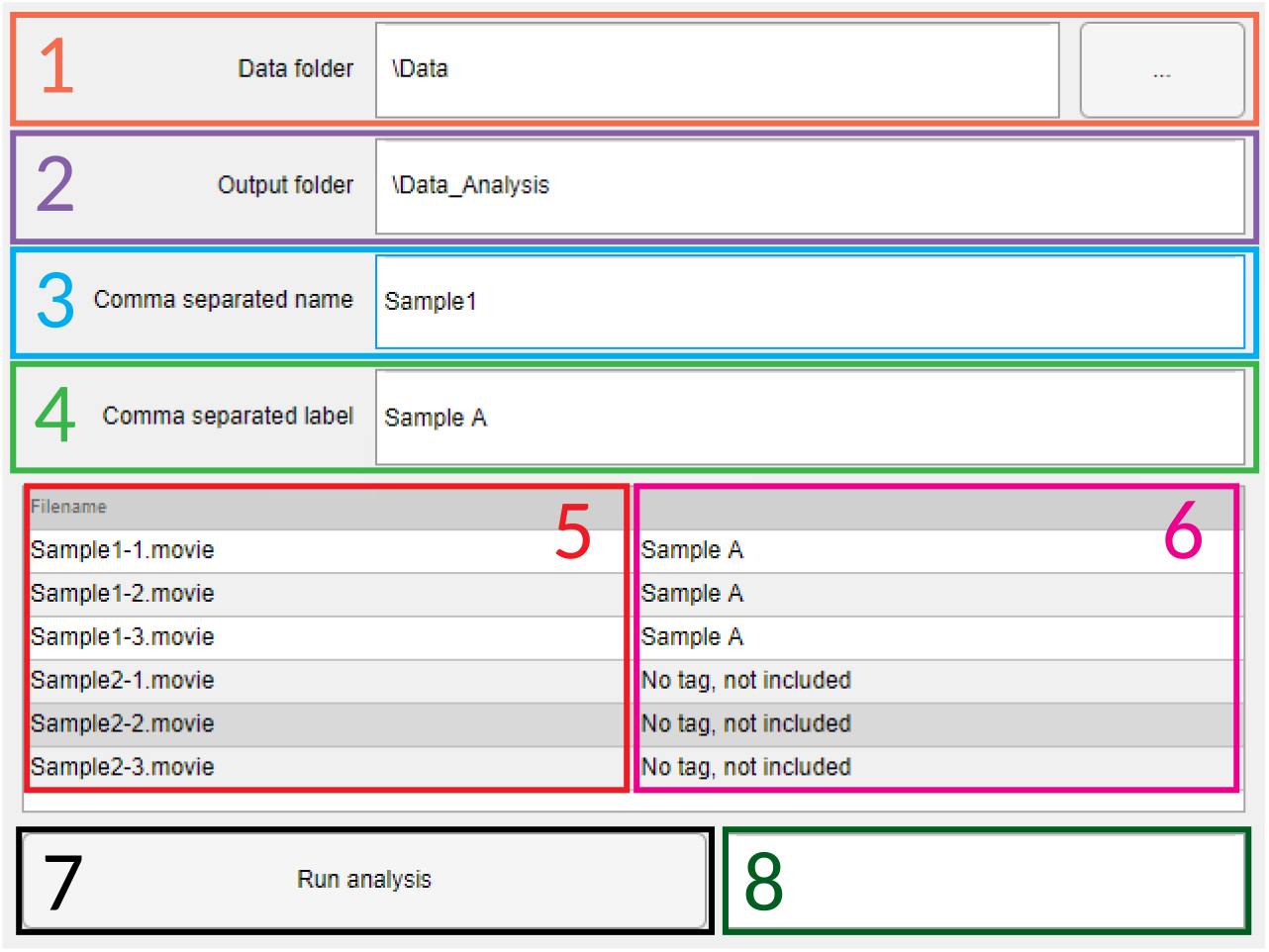
Graphical User Interface used for automatic analysis. 1) Input folder box, the button included can be pressed to search for the correct folder in the filesystem. 2) Output folder box, by definition it will autofill as the input folder plus “_Analysis”. 3) Comma separated list of sample names, this is a list of strings used to distinguish files from the same sample that should be grouped together, coupled with 4) Corresponding list of labels to be used for each sample type in the final plots. 5) List of filenames in the input folder that can be analysed. 6) Information about inclusion for analysis and tag name to be used. 7) Run button, will start the analysis when pressed. 8) Estimated run time based on number of files and time spent per sample

## 5. Results

### (a) Detecting differences in cilia beating frequency and coverage

It is important to be able to clearly distinguish small differences in both beating cilia coverage and their corresponding frequency. Figure 1 shows that this method can easily identify the moving areas in a FOV, from fully covered tissue to very small patches of beating cilia. The algorithm also provide a way to show differences in the CBF distribution for the different videos, in this case displaying a relative decrease in CBF following the decrease in ciliated areas.

### (b) Sample characterization from multiple FOV

In order to measure CBF and coverage across the entire sample, our typical protocol is to collect 20 FOV for each insert, in predefined un-biased positions as described in Figure 8.a. These are selected to include information from both the middle and on the edge areas of the sample, while avoiding any potential selection bias. To estimate a final characterization of a sample, the frequency and movement maps obtained for each FOV are pooled together (Figure 8.b). As described above, the motile cilia percentage is calculated as the accumulated percentage of pixels moving across all the FOV, while the CBF is calculated as the median frequency of all 20 FOV, with the corresponding 25th and 75th percentiles (Figure 8.c).

**Figure 8.**
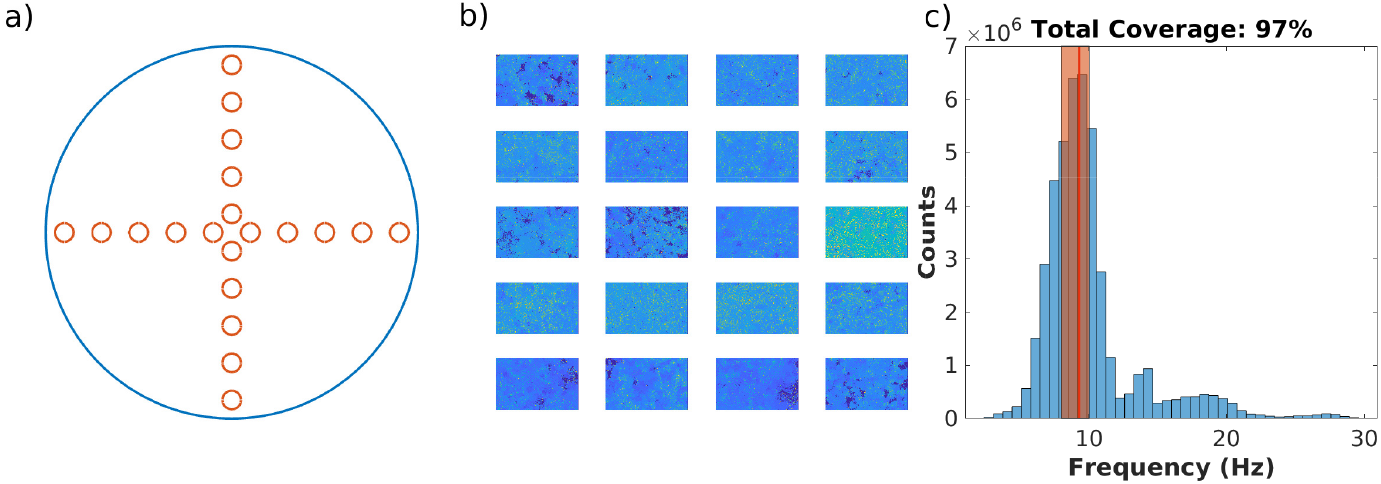
Aggregation method for all FOV collected for a single sample. a) FOV are acquired in a cross shape across the entire insert, including both middle and edge regions. b) All frequency and movement maps for the sample are pooled together. Each FOV is *≈* 187 µm by 117 µm). c) CBF and cilia coverage percentage are calculated based on the aggregated FOV. Red line and orange bar represent the median value, and the 25th-75th percentile interval, respectively.

### (c) Run time and operation

Run time may vary between different systems depending on their hardware configuration. The current version of the algotihm makes use of MATLAB’s GPU acceleration when available. We tested the average run time for the system described in the previous sections, both with GPU acceleration and as CPU only. A FOV movie was first loaded into memory, followed by 5 sequential repeated analysis. While the use of the GPU can drastically decrease the analysis time for each FOV (from about 300 s to about 75 s in this configuration), it is still possible to fully characterise a sample from 20 FOV in about 100 min while only using the CPU (compared with 25 min while using the GPU), highlighting the quick turnaround time for a full sample analysis.

## 6. Conclusion

In this manuscript we presented a new powerful method for the characterization of biophysical properties of airway ciliated epithelium, able to reliably measure CBF and coverage in motile cilia in an automatic fashion. The algorithm we developed is able to independently isolate areas with beating cilia from the background, estimate the distribution of characteristic frequencies in the video, and combine information from multiple collected FOV to characterise the entire sample. The ability to automatically detect motion areas removes a source of human bias in the ROI selection, at the same time decreasing the knowledge barrier required to make use of the software, and increasing the throughput of FOV that can be analysed in a short amount of time. In addition, and unlike most alternatives so far, the ability to quantify and compare the areas of the tissue that presents moving cilia allows for a more thorough characterization of a sample, not only providing information about the frequency at which cilia beat in each specific region, but also measuring how they are distributed across the sample and how this affects their movement. This method also does not require sideways cilia imaging, thus removing further sample processing steps, and is capable of isolating the movement of even small patches of cilia from the background.

## Data Availability

All source code is open-source and available on the repository https://github.com/rgfradique/Canvas Code and sample videos at the time of submission are available here: 10.5281/zenodo.7643501

## Acknowledgment

Funding for this work was from the Cystic Fibrosis Trust grant SRC016 and the EU EC MSCA-ITN grant PhyMot.

